# Loss-functions matter, on optimizing score functions for the estimation of protein models accuracy

**DOI:** 10.1101/651349

**Authors:** Tomer Sidi, Chen Keasar

## Abstract

**Motivation:** Methods for protein structure prediction (PSP) generate multiple alternative structural models (aka decoys). Thus, supervised learning methods for the evaluation and ranking of these models are crucial elements of PSP. Supervised learning involves optimization of loss functions, but their influence on performance is typically overlooked. Here we put the loss functions in the spotlight, and study their effect on prediction performance.

**Results:** Here we report the performances of three variants of MESHI-score, a supervised learning method for the estimation of model accuracy (EMA). Each variant was trained with a different loss function and showed better performance in different aspects of the EMA problem. Most importantly, better discrimination between models of the same target, is gained by target centered loss functions.

**Availability:** All data is available at http://meshi1.cs.bgu.ac.il/SidiAndKeasar2018Data_download/. The MESHI-package (version 9.412) is available at https://github.com/meshiprot/meshi/releases).

## 1 Introduction

Knowledge of protein structures is essential to the understanding of biological processes, and to the design of experimental work in molecular and structural biology. The ultimate sources of this knowledge are experimental methods for structure determination, most notably: X-ray crystallography, NMR and electron microscopy. These methods however are expensive and time consuming, which necessitate complementary computational protein structure prediction (***PSP***) methods.

PSP methods typically generate many alternative structures, henceforth referred to as ***decoys*** (Gront et al., 2012; Jaroszewski et al., 2011; Keasar and Levitt, 2003; Park and Levitt, 1996; Rohl et al., 2004; Söding et al., 2005). Thus, in order to provide the most promising decoys, PSP methods must include some mechanism for the estimation of model accuracy (***EMA*** – aka quality assessment, QA). EMA was recognized as a key aspect of PSP early on (Sippl, 1993), and is still the focus of much research (Elofsson et al., 2017). Since 2008, a dedicated track in the CASP series of PSP experiments measures progress in EMA, and the last rounds have witnessed quite a few impressive results (Kryshtafovych et al., 2017). Yet, the CASP results also indicate that the problem is still open, and none of the current methods is able to consistently pick the best decoy, or accurately predict decoy qualities.

From a methodological perspective, current EMA methods fall into three broad categories: knowledge-based potentials that provide statistical inference from known native structures of proteins (Olechnovič and Venclovas, 2017); unsupervised learning methods that seek consensus within groups of decoys, implicitly assuming random distribution around the native state (McGuffin, 2008; Skwark and Elofsson, 2013; Wallner and Elofsson, 2005); and supervised methods that train on sets of quality-labeled decoys. To this end, supervised methods represent either protein residues or whole proteins as feature vectors and optimize a statistical model to predict protein quality (Manavalan et al., 2014, 2017; Mirzaei et al., 2016; Ray et al., 2012).

The supervised approach is the most general one, as any of the other quality estimate may serve as a feature. Further, it can re-train the statistical model when new sets of labeled decoys become available (*e.g.*, at the end of a CASP experiment). Unfortunately, the adaptation of supervised learning to the EMA problem is none-trivial. Typical supervised learning techniques assume that the data is sampled (at least approximately) from independent and identical distribution (***iid***). EMA techniques cannot assume that, since the EMA problem is inherently group-centered. The estimation of decoy accuracies is meaningful only in the context of their target. Specifically, biologists studying a particular protein, of unknown structure, aim to select the best among decoys of that protein. All these decoys share the same sequence, which distinct them from decoys of other proteins (Figure 1).

**Figure 1.**
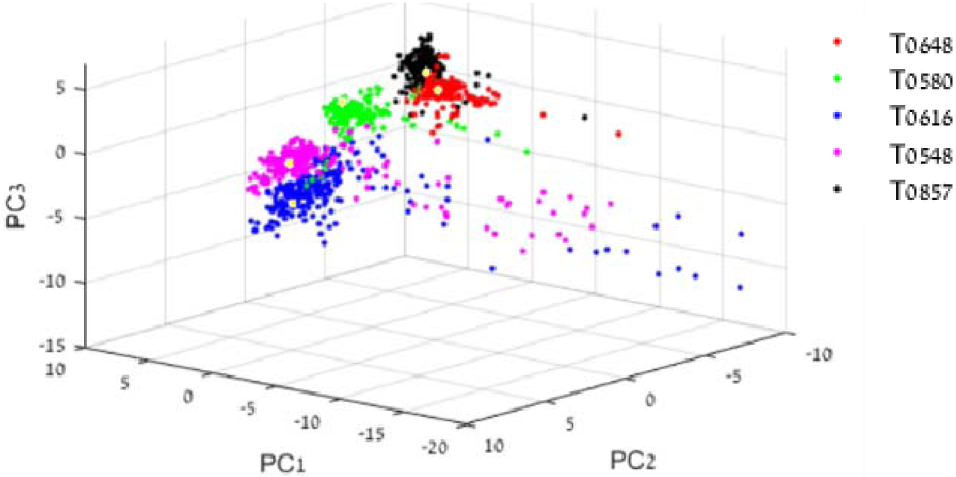
Decoys clusters in feature space. The inherent group-centered nature of EMA may be demonstrated by Principle Component Analysis of feature vectors that represent CASP decoys. For clarity we show only decoys of five arbitrary targets with similar lengths (otherwise the length would have dominated PC1). The Yellow dots depict the best (in terms of GDT_TS) decoy in each set. The decoys of each target occupy an almost distinct region in feature space. High quality decoys are not proximate in this space.

This deviation from ***iid*** distribution of both training and testing data of EMA cannot be ignored. A naïve approach of pooling all data, and blindly applying off-the-shelf tools results in severe overfitting (data not shown). Indeed, all contemporary methods address this challenge by splitting the data into target-disjoint training and test sets, which is compatible with the natural partition of solved and unsolved proteins. The developers of ProQ series of methods (Ray et al., 2012; Uziela et al., 2016) took two steps further: they (a) reduced biases in their training database by randomly choosing the same number of decoys from each target; and (b) they also smoothed the composition differences between targets by applying their machine learning at the residue level, which may be closer to ***iid***.

A major implication of the deviation from ***iid*** in EMA is incompatibility between minimization of the error in accuracy predictions over the entire dataset and per-target measures such as *loss* – the price of missing the best decoy (see below). They coincide only at the apparently unreachable limit of zero error. Yet, methods that use generic supervised learning tools can optimize per-target performance only indirectly by minimizing global error. Qui and his co-workers addressed this difficulty by associating each target of the training set with a binary feature (Larsson et al., 2008). This allowed an arbitrary uniform shift in all the decoy scores of the target during training. Test targets were not associated with such features, resulting in a score that only relate to relative quality. We are not aware of later studies that followed this interesting approach.

MESHI-Score (Elofsson et al., 2017; Keasar et al., 2018; Mirzaei et al., 2016) is a supervised learning method for EMA, that trains its statistical model by Monte-Carlo Simulated Annealing (MCSA) optimization (Brooks and Morgan, 1995; Kirkpatrick, 1984; Metropolis and Ulam, 1949). MCSA is a very flexible computational approach, and is specifically permissive towards the nature of the loss-function that guides optimization. In the context of EMA, it allows either global loss-functions that consider the entire set of decoys, or target-based loss-functions that explicitly consider the group-centric nature of EMA.

Aiming at both absolute and relative accuracy prediction, earlier versions of MESHI-Score used loss-functions that combined a penalty on per-target mean error and a reward for higher Pearson correlation. The utility of the resulted score functions has been demonstrated in CASP11 and CASP12 experiments(Elofsson et al., 2017; Keasar et al., 2018). The current study relaxes the original goal, and seek specialized score functions for either absolute or relative accuracy. Here we report the performances of three MESHI-Score versions. One of them was trained with a global loss-function, and provide the best absolute accuracy estimate. The other two were trained with target-based functions and provide the best relative accuracy estimates. For the sake of completeness, we demonstrate that MESHI-Score is comparable to state-of-the-art EMA method, ProQ3.

## 2 Methods

### 2.1 General

MESHI-Score is an ensemble learning technique for EMA, which aims at predicting the quality of decoy structures (measured in gdt_ts, Zemla, 2003). The overall pipeline of MESHI-score was presented in the previous publications (Elofsson et al., 2017; Mirzaei et al., 2016). In a nutshell, it includes four consecutive steps (Figure 2): (1) decoy standardization; (2) Feature extraction; (3) Parallel prediction by independent predictors; and (4) combining the individual predictions to the ensemble score.

**Figure 2.**
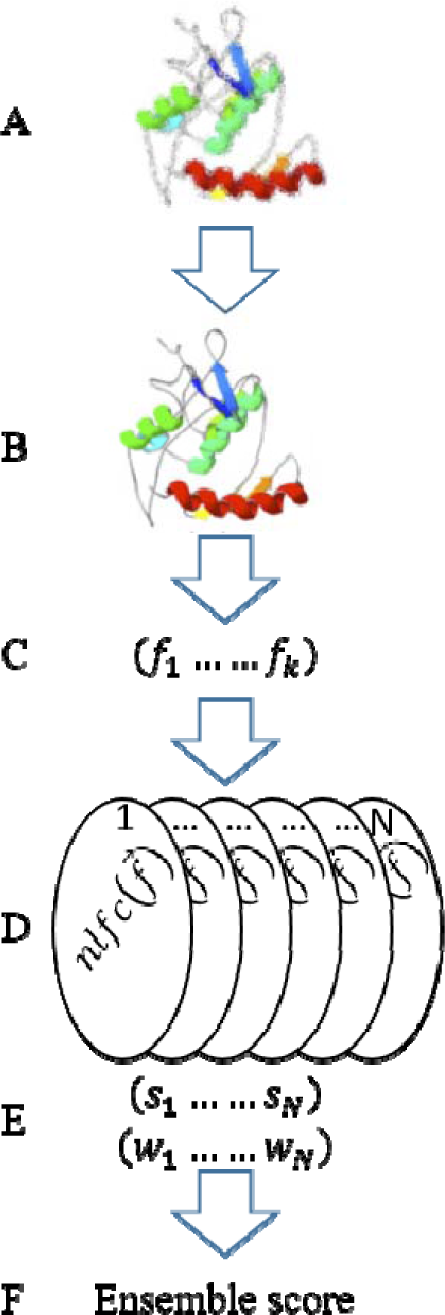
The MESHI-Score pipeline. A. Decoys typically carry the “fingerprints” of the software used for their generation. Specifically, due to small differences between forcefields, very similar models with virtually the same quality may have relatively large distance in feature space. B. The first step in the MESHI-Score pipeline is standardization by sidechain repacking and restrained energy minimization, under the MESHI forcefield. C. Each standardized decoy is represented by K (=129, in the current study) Structural features. D. The feature vector is fed to N (=1000, in the current study) independent predictors. E. Each predictor calculates a different score and weight based on its unique parametrization of the nlfc function. F. The ensemble score is a weighted median of the individual scores.

### 2.2 Database

We extracted and curated a new non-redundant database of decoys from the CASP repository^1^, slightly modifying our previous procedure, by removing the older CASP8 targets, adding newer CASP12 targets and adding a repacking step (see below). The database includes 305 full-chain targets from CASP9-12, with a total of 73605 decoys. Redundant targets, which are homologous to more recent ones, were removed. To this end, we applied all-against-all BLAST (Johnson et al., 2008) comparison to all targets. We considered targets as homologous if either (a) E-value < 10E-6; or (b) E-value is between 10E-6 and 10E-4 and their structures are similar by visual inspection. A standardization procedure reduces structural noise by side-chain repacking using scwrl4 (Krivov et al., 2009) followed by tethered energy minimization (avg. of absolute gdt_ts = 0.007), using the OPTIMIZE program of MESHI-package (version 9.412).

At the end of the tethered minimization step the OPTIMIZE program calculates a vector of 129 features that characterize the decoy. The set of feature vectors constitutes the feature database used to train and test MESHI-score. These features include: (1) pairwise potentials adopted from the literature: GOAP (Zhou and Skolnick, 2011), KB-PMF (Summa and Levitt, 2007), ramp (Samudrala and Moult, 1998), and scwrl 4 energy; (2) in-house developed structural features: torsion angle terms (Amir et al., 2008), quadratic bond and angle terms, radius of gyration, atom environment, and hydrogen bonds (Levy-Moonshine et al., 2009); and (3) compatibility with sequence based prediction of secondary structure and solvent accessibility. These predictions were performed using PSI-PRED (McGuffin et al., 2000) and DeepCNF (Wang et al., 2016), with uniref90 (last updated in 30/7/2017) (Suzek et al., 2015). None of the features considers similarity to other decoys in the set. Full list of the features and descriptions can be found in *https://www.cs.bgu.ac.il/~frankel/TechnicalReports/2015/15-06.pdf*.

### 2.3 Performance measures

All performance measures presented in this manuscript relate to the ability of EMA scores to reproduce decoy accuracies as represented by gdt_ts. Gdt_ts values range between zero (unrelated structures) to one (very similar). Specifically, we use several per-target measures, (Mirzaei et al., 2016), and one global (i.e., applied to all the database decoys):

1. MRMSE - Median per-target root mean square of the prediction error, which is the absolute difference between the decoy’s predicted quality and its gdt_ts score.
2. MC^P^ and MC^S^ - Median per-target Pearson and Spearman Correlation’s respectively, between the predicted and observed qualities.
3. ML - Median per-target loss, the absolute difference between the gdt_ts scores of the top scoring decoy, which is picked by the method, and the actual best decoy in the group.
4. ME10 and ME5 - Median of the per-target enrichment of the and 5% highest accuracy decoys, within the 10% and 5% top scoring ones, respectively. That is, any ME value above one entails better-than-random performance and an ideal predictor reaches ME10 and ME5 values of up to 10 and 20 respectively.
5. GC – Global Spearman correlation between observed and predicted qualities of all decoys

### 2.4 Cross-validation and statistical analysis

To compare the performances of different scores we apply five rounds of 5-fold cross validation. In each round the dataset is randomly partitioned to five sub-sets, each of which serves as the test set once. Overall we get 5X5=25 values per performance measures, and the medians of these values are reported.

While the essence of this manuscript is the comparison of different versions of MESHI-score, we also calculated the EMA score of the state-of-art method ProQ3 (Uziela et al., 2016). ProQ3 is publically-available and we downloaded and ran it using default parameters. Specifically, the ProQ3 values were not generated within the cross validation scheme, and thus their performance is not directly comparable to those of MESHI-score.

### 2.5 Individual predictors

The individual predictors are the only components in MESHI-Score that undergo training. A single predictor is a non-linear combination of features (nlfc - eq.1). To ensure diversity of the predictors, a key ingredient of success in EL (Brown et al., 2005; Sollich and Krogh, 1996), each predictor is associated with a fixed sigmoid function S (eq.2), and is trained to predict the sigmoid transform of decoys’ gdt_ts. Thus, each predictor learns to predict a slightly different objective. To provide a meaningful prediction, the predictor uses the inverse function 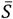 (Eq.3), such that 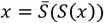. The weight of the prediction, *W* (Eq.4), indicates the sensitivity of the predictor near the predicted value;

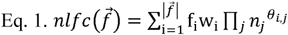

Where 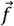 is the features vector, and the normalizers 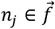 include decoy length, decoy coverage (the fraction of the target sequence that it covers), and the average numbers of 8Å and 15Å Cα contacts of each decoy. w_i_ ∈ ℝ and Θ_*i,j*_ ∈ {1,0.5,0, −0.5, −1, −∞} are the parameters of the statistical model, which are learned from the training data. To reduce the risk of overfitting, no more than fifteen Θ_*i*,1_ values are above -∞, keeping the number of effective features low. Further, for each feature ***i*** only two normalizer exponents (Θ_*i,j*_) are different than zero.

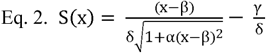

Where 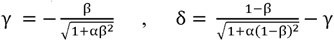, such that 1 ≤ α ≤ 9 and 0 ≤ β ≤ 1 are randomly sampled, with even probability.

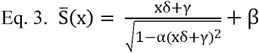

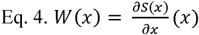

### 2.6 Training of score functions

In MESHI-score, individual predictors learn the parameters of their statistical model through Monte-Carlo Simulated Annealing (MCSA) optimization of a loss function 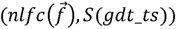. Stochastic optimization methods, like MCSA may optimize any arbitrary loss function, resulting in diverse scores. Here we compare the score functions that resulted from the optimization of three loss functions:

1. *Loss*_*global−score*_= *∝*−*GC* Where GC is global correlation as defined in section 2.3, and ∝ arbitrarily set to one.
2. *Loss*_*corr*_^=^ *β*−*MC* Where MC is per-target median correlation as defined in section 2.3, and *β* arbitrarily set to two.
3. *Loss*_*enrich*10_=1/(1+*Mean*(*Enrichment*10^2^)) Where *Enrichment*10 is per target 10% enrichment.

Given a training set of targets T, and Θ the set of allowed *θ*_*ij*_ combinations (section 2.5), the state of a predictor is (*L, θ*), where *L* ⊂T and *θ* ∈ *Θ*, the weights vector w is derived from the predictors state by averaging the per-target coefficients of linear regression models. The *θ* and *w* vectors are then fed to the score function 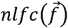 (section 2.5), and the state’s performance, over all the training set T, is evaluated using the loss function. Monte-Carlo steps replace predictor states by a neighboring one (*i.e*, replacing one of *L* ‘s targets).

### 2.7 The ensemble score

In this study, we used ensembles of thousand predictors. Given a feature vector 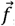, each predictor outputs a pair 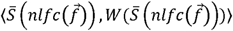. The ensemble score is the weighted median of all the thousand predictions.

## 3 Results and discussion

MESHI-Score trains its individual predictors by MCSA optimization of a loss-function, which provides much flexibility in score development. To study the effect of the loss-function on different performance aspects, we experimented with global and target-based loss functions (data not shown). Here, we systematically compare the three best-performing (to date) versions of MESHI-Score. One of them, MESHI-Score-global-correlation (GC), was trained using a global loss-function (*Loss*_*global−score*_ section 2.7) The other two, MESHI-Score-median-correlation (MC) and MESHI-Score-median-enrichment (ME), were trained using target-centered loss functions (*Loss*_*corr*_ and *Loss*_*enrich*10_ section 2.7). In order to compare the three scores, we performed a 5X5-fold cross validation test. In each round of the test, 4/5 of the targets served as training set, and the three trained scores predicted the decoy accuracies of the remaining fifth. Table 1 presents the average per-target performances of the scores, over six performance measures. Figure 3 provides a more detailed view of the per-target distributions of these performances.

*Tables 1 & 2 – Score performances*

**Table 1.** Each cell is the mean of 5X5 (25 folds) per-target measures. Rows represent the score that were trained using different loss functions (section 2.6). Columns represent the per-target measures as depicted in section 2.3. The best value in each column is bolded. For all measures, the difference between MESHI-Score-global-correlation and the other two scores is statistically significant. (p<10^−2^ Wilcoxon’s signed ranked test for the 25 numbers).

| Table 1 | MC<sup>S</sup> | MC<sup>P</sup> | MRMSE | ME5 | ME10 | ML |
| --- | --- | --- | --- | --- | --- | --- |
| MESHI-Score-global-corr. | 0.6558 | **0.7794** | **0.1198** | 3.7867 | 2.9883 | 0.0617 |
| MESHI-Score-median-corr. | 0.6625 | 0.7292 | 0.1374 | 4.1893 | **3.3250** | **0.0452** |
| MESHI-Score-median-enric. | **0.6695** | 0.7387 | 0.1367 | **4.2500** | **3.3250** | 0.0493 |

**Figure 3.**
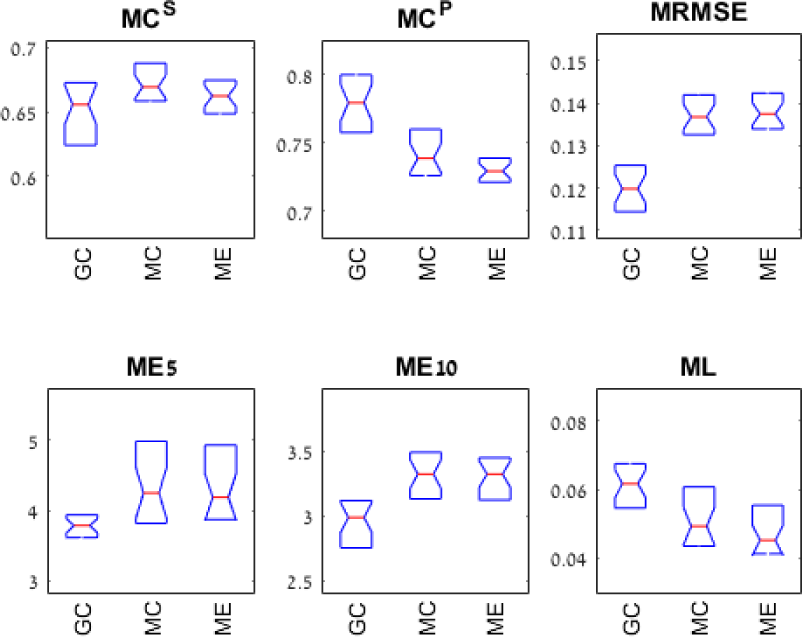
Per-target distribution of the performances. GC, MC, and ME refer to the three score functions: MESHI-Score-global-correlation, MESHI-Score-median-correlation, and MESHI-Score-median-enrichment respectively. MC^S^, MC^P^, MRMSE, ME5, ME10, and ML refer to median: Spearman correlation, Pearson correlation, root mean square of error, 5% enrichment, 10% enrichment, and loss, respectively. The two segments of each box indicate the second and third quantiles of the population

Overall the three scores have similar performances, which are on-par with the current state of the art (Table 2). Yet, the differences between the global score and the two target-centered scores are statistically significant and telling. Most notably, the different measures are not consistent with one another. The best prediction of mean absolute accuracy, is achieved by the global score (GC), yet it performs worse in estimating the relative accuracies of decoys within their group. The target-centered scores on the other hand are more likely to pick one of the best decoy available, and less likely to pick one of the least accurate decoys (Figure 4 and Figure 5).

**Table 2.** ProQ3 - Median of per-target measures, with default parameters. Note that data for 758 decoys are missing due to software failures. The results of ProQ3 cannot be quantitatively compared with those of MESHI-Score as the latter were not created within the cross validation scheme.

|  |  |  |  |  |  |  |
| --- | --- | --- | --- | --- | --- | --- |
| ProQ3 | 0.6138 | 0.7044 | 0.1794 | 3.7200 | 2.8666 | 0.0581 |

**Figure 4.**
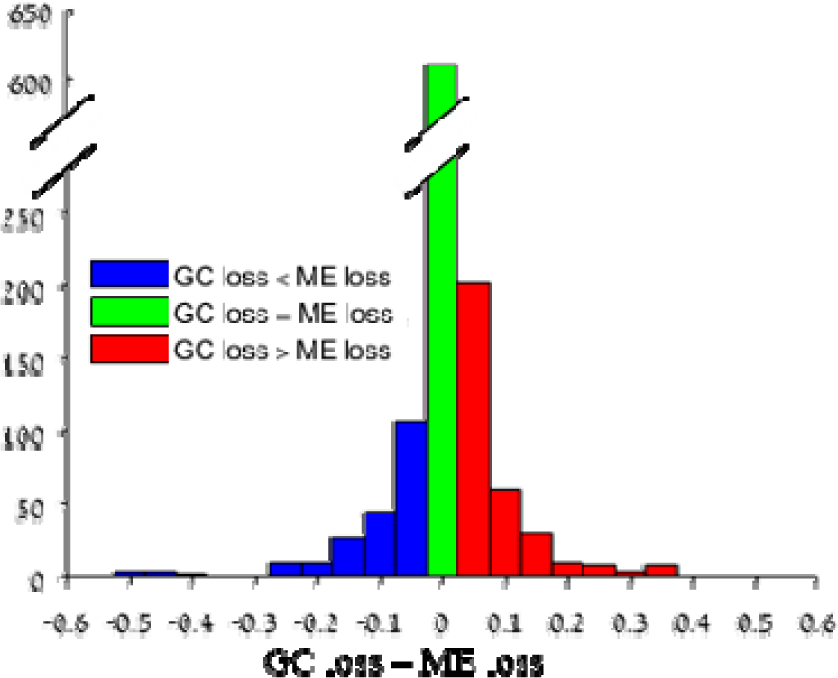
The distribution of differences in loss values between MESHI-Score-median-enrichment (ME) and MESHI-Score-global-correlation (GC). Five GC/ME pairs are generated by *5X5 cross-validation test for each database target. The histogram depicts the distribution of their loss value differences. The apparent skew towards positive (red) differences indicates that ME is more likely to result lower losses than GC. Similar trends are observed also for the other measures of Table 1, and Wilcoxon signed rank test indicate that their skews are statistically significant (Wilcoxon signed rank test; P<0.01)*.

**Figure 5.**
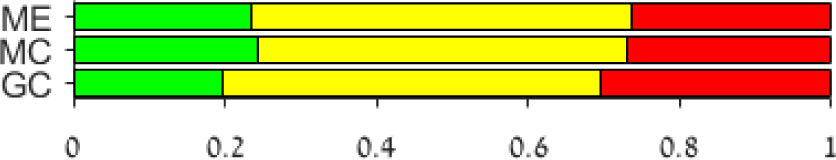
Success and failure rates in picking the best models available. Fractions of targets for which high quality decoys were picked by the score function (green = loss 0.02), 0.02 loss 0.1, and loss 0.1, Green, orange, and red, respectively

## 4 Conclusions

The EMA problem is a long standing hard computational problem, which gets complicated by its group-centered nature. Further, it is a multi-objective problem as we seek both absolute and relative estimates of accuracy. Our results suggest that the loss-function, which is used to optimize the parameters of an EMA score, has a profound impact on the score’s performance, and that different loss-functions may excel in different objectives. Specifically, target-based loss-functions may be beneficial for target-based objectives (*e.g.*, loss). One may speculate that this insight may be applicable to other machine-learning tasks, especially ones in which the standard assumption of ***iid*** sampling of the data is not applicable.

## Acknowledgements

The authors are grateful to the CASP server groups and to the Prediction Center for generating, archiving and providing all the data used in this study.

## Funding

The authors are grateful for support by grant no. 1122/14 from the Israel Science Foundation (ISF).

## Conflict of Interest

none declared.

## Footnotes

1 http://predictioncenter.org

## References

Amir, E.-A.D., Kalisman, N., and Keasar, C. (2008). Differentiable, multidimensional, knowledge-based energy terms for torsion angle probabilities and propensities. Proteins Struct. Funct. Bioinformaxs. 72, 62–73.

Brooks, S.P., and Morgan, B.J.T. (1995). Optimization Using Simulated Annealing. J. R. Stat. Soc. Ser. Stat. 44, 241–257.

Brown, G., Wyatt, J., Harris, R., and Yao, X. (2005). Diversity creation methods: a survey and categorisation. Inf. Fusion 6, 5–20.

Elofsson, A., Joo, K., Keasar, C., Lee, J., Maghrabi, A.H.A., Manavalan, B., McGuffin, L.J., Hurtado, D.M., Mirabello, C., Pilstål, R., et al. (2017). Methods for estimation of model accuracy in CASP12. Proteins Struct. Funct. Bioinforma. 86, 361–373.

Gront, D., Kmiecik, S., Blaszczyk, M., Ekonomiuk, D., and Koliński, A. (2012). Optimization of protein models. Wiley Interdiscip. Rev. Comput. Mol. Sci. 2, 479–493.

Jaroszewski, L., Li, Z., Cai, X., Weber, C., and Godzik, A. (2011). FFAS server: novel features and applications. Nucleic Acids Res. 39, W38–W44.

Johnson, M., Zaretskaya, I., Raytselis, Y., Merezhuk, Y., McGinnis, S., and Madden, T.L. (2008). NCBI BLAST: a better web interface. Nucleic Acids Res. 36, W5–W9.

Keasar, C., and Levitt, M. (2003). A novel approach to decoy set generation: designing a physical energy function having local minima with native structure characteristics., A Novel Approach to Decoy Set Generation: Designing a Physical Energy Function Having Local Minima with Native Structure Characteristics. J. Mol. Biol. J. Mol. Biol. 329, 329, 159, 159–174.

Keasar, C., McGuffin, L.J., Wallner, B., Chopra, G., Adhikari, B., Bhattacharya, D., Blake, L., Bortot, L.O., Cao, R., Dhanasekaran, B.K., et al. (2018). An analysis and evaluation of the WeFold collaborative for protein structure prediction and its pipelines in CASP11 and CASP12. Sci. Rep. 8, 9939.

Kirkpatrick, S. (1984). Optimization by simulated annealing: Quantitative studies. J. Stat. Phys. 34, 975–986.

Krivov, G.G., Shapovalov, M.V., and Dunbrack, R.L. (2009). Improved prediction of protein side-chain conformations with SCWRL4. Proteins Struct. Funct. Bioinforma. 77, 778–795.

Kryshtafovych, A., Monastyrskyy, B., Fidelis, K., Schwede, T., and Tramontano, A. (2017). Assessment of model accuracy estimations in CASP12. Proteins Struct. Funct. Bioinforma. 86, 345–360.

Larsson, P., Wallner, B., Lindahl, E., and Elofsson, A. (2008). Using multiple templates to improve quality of homology models in automated homology modeling. Protein Sci. 17, 990–1002.

Levy-Moonshine, A., Amir, E.D., and Keasar, C. (2009). Enhancement of betasheet assembly by cooperative hydrogen bonds potential. Bioinformatics 25, 2639–2645.

Manavalan, B., Lee, J., and Lee, J. (2014). Random Forest-Based Protein Model Quality Assessment (RFMQA) Using Structural Features and Potential Energy Terms. PLOS ONE 9, e106542.

Manavalan, B., Lee, J., and Valencia, A. (2017). SVMQA: support–vector-machine-based protein single-model quality assessment. Bioinformatics 33, 2496–2503.

McGuffin, L.J. (2008). The ModFOLD server for the quality assessment of protein structural models. Bioinformatics 24, 586–587.

McGuffin, L.J., Bryson, K., and Jones, D.T. (2000). The PSIPRED protein structure prediction server. Bioinformatics 16, 404–405.

Metropolis, N., and Ulam, S. (1949). The Monte Carlo Method. J. Am. Stat. Assoc. 44, 335–341.

Olechnovič, K., and Venclovas, Č. (2017). VoroMQA: Assessment of protein structure quality using interatomic contact areas. Proteins Struct. Funct. Bioinforma. 85, 1131–1145.

Park, B., and Levitt, M. (1996). Energy Functions that Discriminate X-ray and Near-native Folds from Well-constructed Decoys. J. Mol. Biol. 258, 367–392.

Ray, A., Lindahl, E., and Wallner, B. (2012). Improved model quality assessment using ProQ2. BMC Bioinformatics 13, 224.

Rohl, C.A., Strauss, C.E.M., Misura, K.M.S., and Baker, D. (2004). Protein Structure Prediction Using Rosetta. In Methods in Enzymology, (Academic Press), pp. 66–93.

Samudrala, R., and Moult, J. (1998). An all-atom distance-dependent conditional probability discriminatory function for protein structure prediction11Edited by F. Cohen. J. Mol. Biol. 275, 895–916.

Sippl, M.J. (1993). Recognition of errors in three-dimensional structures of proteins. Proteins Struct. Funct. Bioinforma. 17, 355–362.

Skwark, M.J., and Elofsson, A. (2013). PconsD: ultra rapid, accurate model quality assessment for protein structure prediction. Bioinformatics 29, 1817–1818.

Söding, J., Biegert, A., and Lupas, A.N. (2005). The HHpred interactive server for protein homology detection and structure prediction. Nucleic Acids Res. 33, W244–W248.

Sollich, P., and Krogh, A. (1996). Learning with ensembles: How overfitting can be useful. In Advances in Neural Information Processing Systems 8, D.S. Touretzky, and M.E. Hasselmo, eds. (MIT Press), pp. 190–196.

Summa, C.M., and Levitt, M. (2007). Near-native structure refinement using in vacuo energy minimization. Proc. Natl. Acad. Sci. 104, 3177–3182.

Suzek, B.E., Wang, Y., Huang, H., McGarvey, P.B., and Wu, C.H. (2015). UniRef clusters: a comprehensive and scalable alternative for improving sequence similarity searches. Bioinformatics 31, 926–932.

Uziela, K., Shu, N., Wallner, B., and Elofsson, A. (2016). ProQ3: Improved model quality assessments using Rosetta energy terms. Sci. Rep. 6, 33509.

Wallner, B., and Elofsson, A. (2005). Pcons5: combining consensus, structural evaluation and fold recognition scores. Bioinformatics 21, 4248–4254.

Wang, S., Li, W., Liu, S., and Xu, J. (2016). RaptorX-Property: a web server for protein structure property prediction. Nucleic Acids Res. 44, W430–W435.

Zemla, A. (2003). LGA: a method for finding 3D similarities in protein structures. Nucleic Acids Res. 31, 3370–3374.

Zhou, H., and Skolnick, J. (2011). GOAP: A Generalized Orientation-Dependent, All-Atom Statistical Potential for Protein Structure Prediction. Biophys. J. 101, 2043–2052.

